# Risk factors for asthma among schoolchildren who participated in a case-control study in urban Uganda

**DOI:** 10.1101/677799

**Authors:** Harriet Mpairwe, Milly Namutebi, Gyaviira Nkurunungi, Pius Tumwesige, Irene Nambuya, Mike Mukasa, Caroline Onen, Marble Nnaluwooza, Barbara Apule, Tonny Katongole, Gloria Oduru, Joseph Kahwa, Emily L Webb, Lawrence Lubyayi, Neil Pearce, Alison M Elliott

**Affiliations:** Medical Research Council/Uganda Virus Research Institute and London School of Hygiene and Tropical Medicine Uganda Research Unit. Plot 51-59 Nakiwogo Road, Box 49, Entebbe, Uganda; London School of Hygiene and Tropical Medicine. Keppel St, Bloomsbury, London WC1E 7HT, UK

**Author notes:** Corresponding author: Harriet Mpairwe.

**Keywords:** childhood asthma, case-control, allergic sensitisation, early life, rural, Uganda

## Abstract

Data on asthma aetiology in Africa are scarce. We investigated the risk factors for asthma among schoolchildren (5-17years) in urban Uganda. We conducted a case-control study, enrolling 561 cases and 1,139 controls. Asthma was diagnosed by study clinicians.

The main risk factors for asthma were tertiary education for fathers [adjusted OR (95% CI); 2.49 (1.89-3.29)] and mothers [2.14 (1.64-2.78)]; area of residence at birth, with children born in a small town or in the city having an increased asthma risk compared to schoolchildren born in rural areas [2.00 (1.49-2.68)] and [2.82 (1.85-4.30)], respectively; father’s and mother’s history of asthma; children’s own allergic conditions; atopy; and using gas/electricity for indoor cooking.

Asthma was associated with a strong rural-town-city risk gradient, higher parental socio-economic status and urbanicity. This work provides the basis for future studies to identify specific environmental/lifestyle factors responsible for increasing asthma risk among children in urban areas in LMICs.

## Background

Asthma is estimated to affect more than 235 million people globally, and is the most common non-communicable condition among children(1). In Africa, the prevalence of asthma appears to be increasing(2–6), particularly in urban areas(3, 7), but the causes of this increase are not fully understood. Moreover, asthma has various phenotypes which may have different aetiologies(8), and risk factors appear to vary internationally, and to differ between high-income countries (HICs) and low-and-middle income countries (LMICs)(9).

There is a scarcity of data on asthma risk factors from Africa. The few studies reported have suggested that current residence in urban areas is associated with a higher risk of asthma than rural residence(10, 11). Other risk factors, similar to those in HICs, include maternal smoking(12, 13), maternal history of asthma(14), childhood atopic sensitisation(11, 15) and history of allergy(14, 16). Previous reports suggest no association between biomass fuels and asthma risk(17, 18), but increased asthma symptoms(18, 19).

We undertook a case-control study among schoolchildren in an urban area in Uganda, to investigate the main risk factors for asthma and the patterns of allergic sensitisation.

## Methods

### Study design

We conducted a case-control study among schoolchildren, and report following STROBE guidelines(20).

### Study population, sampling and consent

The study was conducted in 5-17-year-old schoolchildren in primary and secondary schools in Wakiso District, Central Uganda, a predominantly urban setting.

All schools in the study area were invited and 96% participated. At each school, we prescreened children by requesting all those with any breathing problems to register with us, and concurrently recruited two children without any breathing problems at random from the same class, using a random number generator programme in STATA (StataCorp, Texas, USA). The children delivered invitation letters to their parents; parents/guardians with telephones were invited to attend a meeting during which those interested provided written informed consent, and children aged ≥8 years provided written informed assent. Study enrollment was between May 2015 and July 2017.

### Ethical approval

The study was approved by the Uganda Virus Research Institute Research and Ethics Committee, and the Uganda National Council for Science and Technology.

### Definition of cases and controls

Children with a history of wheezing in the last 12 months according to the International Study of Asthma and Allergies in Childhood (ISAAC) questionnaire (21) underwent a detailed medical history and examination by study clinicians, to diagnose asthma; if the diagnosis was not straightforward, two clinicians reviewed that participant and if they disagreed, that participant was excluded. Children with a history of wheeze or any asthma symptoms but not in the last 12 months were excluded. Controls were defined as having no history of wheeze or any other asthma symptoms. There were no other exclusion criteria.

### Clinical assessments

We collected data about asthma risk factors and allergic conditions using interviewer-led questionnaires to parents and adolescents, including the ISAAC questionnaire (21). When a parent was not available in person, we interviewed them by telephone. We examined for presence of a Bacillus Calmette–Guérin (BCG) scar, and tested for fractional exhaled nitric oxide (FENO), using a hand-held device (NoBreath^®^ from Bedfonf Scientific, Maidstone, United Kingdom). We used the manufacturer’s cut-off for children of ≥35 parts per billion.

We conducted skin prick tests (SPT) following standard procedures(22), and crude extracts of seven allergens (*Dermatophagoides* mix of *D. farinae* and *D. pteronyssminus*, *Blomia tropicalis*, *Blattella germanica*, peanut, cat, pollen mix of weeds, mould mix of *Aspergillus* species; ALK Abello, Hoersholm, Denmark). A positive response was a wheal diameter ≥3mm measured after 15 minutes, with a negative saline control and positive histamine. We collected blood samples which we processed to obtain plasma that we stored at −80ºC. At the end of the study, we randomly selected 200 plasma aliquots from all asthma cases and 200 from all controls for measurement of allergen-specific IgE (asIgE) to whole allergen extracts (*D. pteronyssinus*, *B. germanica* and peanut), using ImmunoCAP^®^ (Phadia, Uppsala, Sweden). The standard cut-off for allergic sensitisation of ≥0.35 allergen-specific kilo units per litre (kU_A_/L) was used.

We also collected three fresh stool samples from each participant to test for intestinal helminths using the Kato Katz method(23). We performed the tuberculin skin test (TST) using standard procedures as we have previously described(24).

### Data management and statistical analysis

Data were double-entered using OpenClinica open source software version 3.1.4 (OpenClinica LLC and collaborators, Waltham, MA, USA). Data were analysed in STATA. We estimated that a sample size of 2,112 would have 80% power to detect odds ratio (OR) <0.5 or >1.5 for exposures with prevalence 10% among controls.

For continuous variables with clinically relevant cut-off points such as for SPT, asIgE, FENO and TST, we used the dichotomous variables in the analysis. Total IgE was analysed as a continuous variable. The variable for maternal or paternal history of ‘allergic disease’ combines the history of asthma, eczema, allergic rhinitis, allergic conjunctivitis and any other allergies. We conducted a complete case analysis, and did not impute missing values.

Controls were not matched to cases by age or sex; rather, they were randomly sampled from the ‘non-cases’ as described above, and we then controlled for age and sex in the analysis. We conducted initial logistic regression analyses for each exposure of interest, adjusted for age and sex. We built the final multiple logistic regression models by adding one potential confounder (identified in the literature) at a time and examined the change in the main effect estimate, to obtain adjusted ORs, and stopped adding variables when the effect estimate stopped changing(25, 26). Factors on the causal pathway were not adjusted for, and closely related factors (such as father’s and mother’s education level, or area of residence at the time of birth and area where the child spent 0-5 years) were not included in the same model, to avoid multicollinearity(26).

We assessed whether the combined effect of having two of the main asthma risk factors was additive or multiplicative.

## Results

### Reference characteristics of participating schools and participant flow

We enrolled participants from 55 schools (32 primary, 23 secondary). Of the 6,385 children initially identified from the pre-screening exercise, we were unable to contact about half of the parents/guardians in time for them to attend the parents’ meeting; most of these children were in the boarding section and their homes were outside the study area (Figure 1). Of the 1,835 who attended the meeting, 97% provided written informed consent for their child to participate in the study. We screened 1,779 participants and of these, 77 who had initially reported breathing problems either did not have an asthma diagnosis or did not have asthma symptoms in the last 12 months and were excluded. We enrolled 562 children with and 1,140 without asthma, but excluded two with incomplete data (Figure 1).

**Figure 1:**
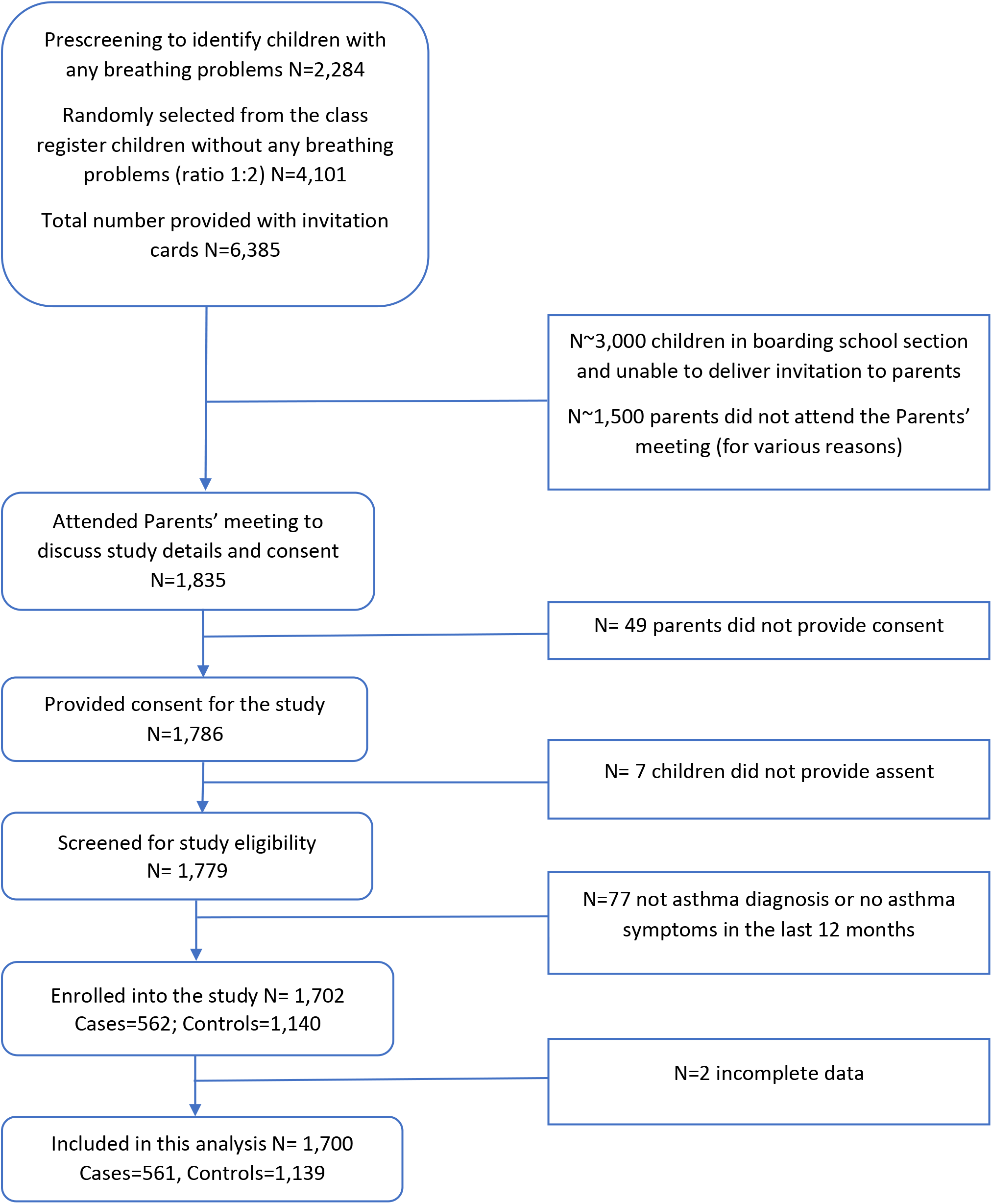
Participant flow diagram.

### Early life risk factors for asthma

Participants had mean age 11 years (range 5-17 years); children with asthma were slightly older, and more likely to have parents with a tertiary education and a reported history of asthma (Table 1). Compared to children born in rural Uganda, children born in any town in Uganda or in the city had an increased risk of asthma [adjusted OR (95% confidence interval (CI)) 2.00 (1.49-2.68)] and [2.82 (1.85-4.30)], respectively. The same pattern was observed for the area where the child spent most of their early life (0-5 years) (Table 1). There were no differences in reported exposure to farm animals, or to cigarette smoke during pregnancy. Children with asthma were less likely to have a BCG scar [0.67 (0.51-0.88), p-value=0.004] (Table 1), but the TST response (induration ≥10mm) at enrolment was similar between cases (15.7%) and controls (14.6%) [1.01 (0.68-1.50)]. There was no statistical evidence of interaction between parental education and the children’s area of residence in early life, nor interaction between age and any of the asthma risk factors.

**Table 1:** Risk factors in early life among asthma cases and non-asthma controls.

| **Risk factors** | **Asthma cases**<br>**N=561** | **Non-asthma controls**<br>**N=1,139** | **Adj. OR<sup>‡</sup> (95% CI)** | **P-value** |
| --- | --- | --- | --- | --- |
| **Age**, years, Mean (Range) [m=3] | 11.4 (5-17) | 10.9 (5-17) | 1.08 (1.04-1.11) | <0.0001 |
| **Sex:** Girls (944) | 296 (52.8) | 648 (56.9) | 0.82 (0.66-1.01) | 0.06 |
| **Mother's history of asthma** [m=181] |  |  |  |  |
| Yes (103) | 61 (12.0) | 42 (4.2) | 3.05 (1.99-4.66) | <0.0001 |
| **Father's history of asthma** [m=196] |  |  |  |  |
| Yes (85) | 65 (12.8) | 20 (2.0) | 6.61 (3.92-11.15) | <0.0001 |
| **Fathers' highest education level** [m=29] |  |  |  |  |
| None/Primary (437) | 99 (17.8) | 338 (30.3) | 1 |  |
| Secondary (595) | 194 (34.9) | 401 (36.0) | 1.56 (1.17-2.08) | <0.0001 |
| Tertiary (639) | 263 (47.3) | 376 (33.7) | 2.22 (1.67-2.94)<sup>#</sup> |  |
| **Mothers' highest education attained** [m=30] |  |  |  |  |
| None/Primary (580) | 159 (28.7) | 421 (37.7) | 1 |  |
| Secondary (602) | 185 (33.4) | 417 (37.4) | 1.10 (0.85-1.42) | <0.0001 |
| Tertiary (488) | 210 (37.9) | 278 (24.9) | 1.85 (1.41-2.42)<sup>#</sup> |  |
| **Area of residence at the time of the child's birth** [m=8] |  |  |  |  |
| Rural (355) | 75 (13.4) | 280 (24.7) | 1 |  |
| Town (1,188) | 416 (74.3) | 772 (68.2) | 2.00 (1.49-2.68) | <0.0001 |
| City (149) | 69 (12.3) | 80 (7.1) | 2.82 (1.85-4.30)<sup>#</sup> |  |
| **Area of residence where the child spent most of 0-5 years** [m=8] |  |  |  |  |
| Rural (339) | 72 (12.9) | 267 (23.6) | 1 |  |
| Town (1,267) | 439 (78.4) | 828 (73.1) | 1.84 (1.37-2.48) | <0.0001 |
| City (86) | 49 (8.7) | 37 (3.3) | 3.77 (2.25-6.30)<sup>#</sup> |  |
| **Maternal regular contact with farm animals during pregnancy** [m=270] |  |  |  |  |
| Yes (504) | 152 (30.2) | 352 (38.0) | 0.83 (0.65-1.05) | 0.12 |
| **Maternal cigarette smoking during pregnancy** [m=320] |  |  |  |  |
| Yes (36) | 13 (2.6) | 23 (2.6) | 1.11 (0.54-2.67) | 0.77 |
| **Breast feeding duration** [m=420] |  |  |  |  |
| ≤1year (333) | 114 (25.22) | 219 (26.45) | 1 |  |
| >1year (947) | 338 (74.78) | 609 (73.55) | 1.10 (0.84-1.45) | 0.47 |
| **Allergic rhinitis ever** [m=8] |  |  |  |  |
| Yes (870) | 431 (77.1) | 439 (38.8) | 4.90 (3.77-6.38) | <0.0001 |
| **Allergic conjunctivitis ever** [m=9] |  |  |  |  |
| Yes (772) | 364 (64.1) | 408 (36.0) | 2.87 (2.26-3.65) | <0.0001 |
| **Eczema ever** [m=9] |  |  |  |  |
| Yes (265) | 126 (22.5) | 139 (12.3) | 1.84 (1.35-2.50) | <0.0001 |
| **Urticarial rash ever** [m=9] |  |  |  |  |
| Yes (572) | 223 (39.9) | 349 (30.8) | 1.45 (1.13-1.86) | 0.003 |
| **Presence of a BCG scar on child's arm at enrolment** [m=90] |  |  |  |  |
| Yes (1,335) | 417 (78.4) | 918 (85.2) | 0.67 (0.51-0.88) | 0.004 |

Adj. OR=adjusted odds ratio; CI=confidence interval; N=number; m=missing; number (%) shown in 2<sup>nd</sup> and 3<sup>rd</sup> column unless indicated otherwise. <sup>‡</sup>adjusted for child's age, sex, area of residence at the time of birth, and father's education (additionally, children's allergic disease was adjusted for father's and mother's history of asthma). <sup>#</sup>Test for trend p-value <0.0001

### Current features of asthma cases versus controls

Asthma cases were more likely to report a high frequency of ‘trucks passing on the street near their home’ [2.35 (1.59-3.48)]; to come from homes that used electricity/gas for indoor cooking [1.56 (1.15-2.11)]; and to report having used de-worming medication more than twice in the last 12 months [2.32 (1.75-3.07)] than controls. There was weak evidence for an inverse association between asthma and infection with any helminths species [0.72 (0.51-1.02)] (Table 2), and this was significant for *T. trichiura* [0.32 (0.12-0.85)] (Supplementary Table 1). Children with asthma were more likely to report a history of allergic diseases such as allergic rhinitis, conjunctivitis, eczema and urticarial rashes (Table 1) and to have these conditions currently (Table 2). The prevalence of current exposure to cigarette smoke (in the household) was similar among cases and controls, although only about 10% were exposed (Table 2).

**Table 2:** Current features of asthma cases and non-asthma controls.

| **Current features** | **Asthma cases**<br>**N=561** | **Non-asthma controls**<br>**N=1,139** | **Adj. OR\* (95% CI)** | **P-value** |
| --- | --- | --- | --- | --- |
| **Frequency of 'Trucks' passing on street near child's home [m=8]** |  |  |  |  |
| Rarely (945) | 269 (48.0) | 676 (59.7) | 1 |  |
| Frequently (624) | 232 (41.4) | 392 (34.6) | 1.59 (1.27-1.99) | <0.0001 |
| Almost all the time (123) | 59 (10.5) | 64 (5.7) | 2.21 (1.48-3.28)<sup>#</sup> |  |
| **Cooking fuel most frequently used indoor [m=8]** |  |  |  |  |
| No indoor cooking (394) | 121 (21.6) | 273 (24.1) | 1 |  |
| Charcoal stove (908) | 266 (47.5) | 642 (56.7) | 1.02 (0.78-1.33) | 0.88 |
| Gas/Electricity (390) | 173 (30.9) | 217 (19.2) | 1.55 (1.15-2.10) | 0.005 |
| **Current exposure to cigarette smoke by a household member [m=7]** |  |  |  |  |
| Yes (191) | 70 (12.5) | 121 (10.7) | 1.31 (0.94-1.82) | 0.11 |
| **Use of de-worming medication in last 12 months [m=10]** |  |  |  |  |
| None (578) | 151 (27.0) | 427 (37.8) | 1 |  |
| Once (732) | 232 (41.5) | 500 (44.2) | 1.36 (1.06-1.74) | <0.0001 |
| ≥Twice (380) | 176 (31.5) | 204 (18.0) | 2.19 (1.64-2.90)<sup>#</sup> |  |
| **Used albendazole in last 12 months [m=27]** |  |  |  |  |
| Yes (905) | 355 (64.0) | 550 (49.2) | 1.75 (1.41-2.18) | <0.0001 |
| **Used praziquantel in last 12 months [m=64]** |  |  |  |  |
| Yes (401) | 116 (21.4) | 285 (26.1) | 0.86 (0.66-1.11) | 0.24 |
| **Presence of any helminth infection at enrolment<sup>†</sup> [m=134]** |  |  |  |  |
| Yes (213) | 55 (10.6) | 158 (15.1) | 0.72 (0.51-1.02) | 0.06 |
| **Allergic rhinitis in last 12 months [m=9]** |  |  |  |  |
| Yes (216) | 119 (21.3) | 97 (8.6) | 2.81 (1.99-3.96) | <0.0001 |
| **Allergic conjunctivitis in last 12 months [m=9]** |  |  |  |  |
| Yes (179) | 86 (15.4) | 93 (8.2) | 1.85 (1.28-2.68) | 0.001 |
| **Eczema in last 12 months [m=9]** |  |  |  |  |
| Yes (60) | 41 (7.3) | 19 (1.7) | 4.48 (2.37-8.49) | <0.0001 |
| **Urticarial rash in last 12 months [m=9]** |  |  |  |  |
| Yes (36) | 18 (3.2) | 18 (1.6) | 2.80 (1.15-6.82) | 0.02 |

Adj. OR=adjusted odds ratio; CI=confidence interval; N=number; m=missing; number (%) shown in 2<sup>nd</sup> and 3<sup>rd</sup> column unless indicated otherwise. \*adjusted for child's age, sex, area of residence at the time of birth, and father's education (additionally, children's allergic disease was adjusted for father's and mother's history of asthma). <sup>#</sup>Test for trend p-value <0.0001. <sup>†</sup>Helminths infections included mainly *Schistosoma mansoni*, *Trichuris trichuria*, Hookworm, and *Ascaris lumbricoides*; this variable was additionally adjusted for reported worm treatment in the last 12 months.

### Atopy and asthma

Children with asthma were more likely to have allergic sensitisation: SPT positive to at least one of seven whole allergen extracts [2.46 (1.98-3.05)]; and elevated asIgE levels to any of three whole allergen extracts [2.35 (1.51-3.66)], and higher total IgE (Table 3). The most common allergens were dust mites and cockroach. Asthma cases were more likely to have elevated FENO levels [2.63 (2.06-3.35)] (Table 3).

**Table 3:**
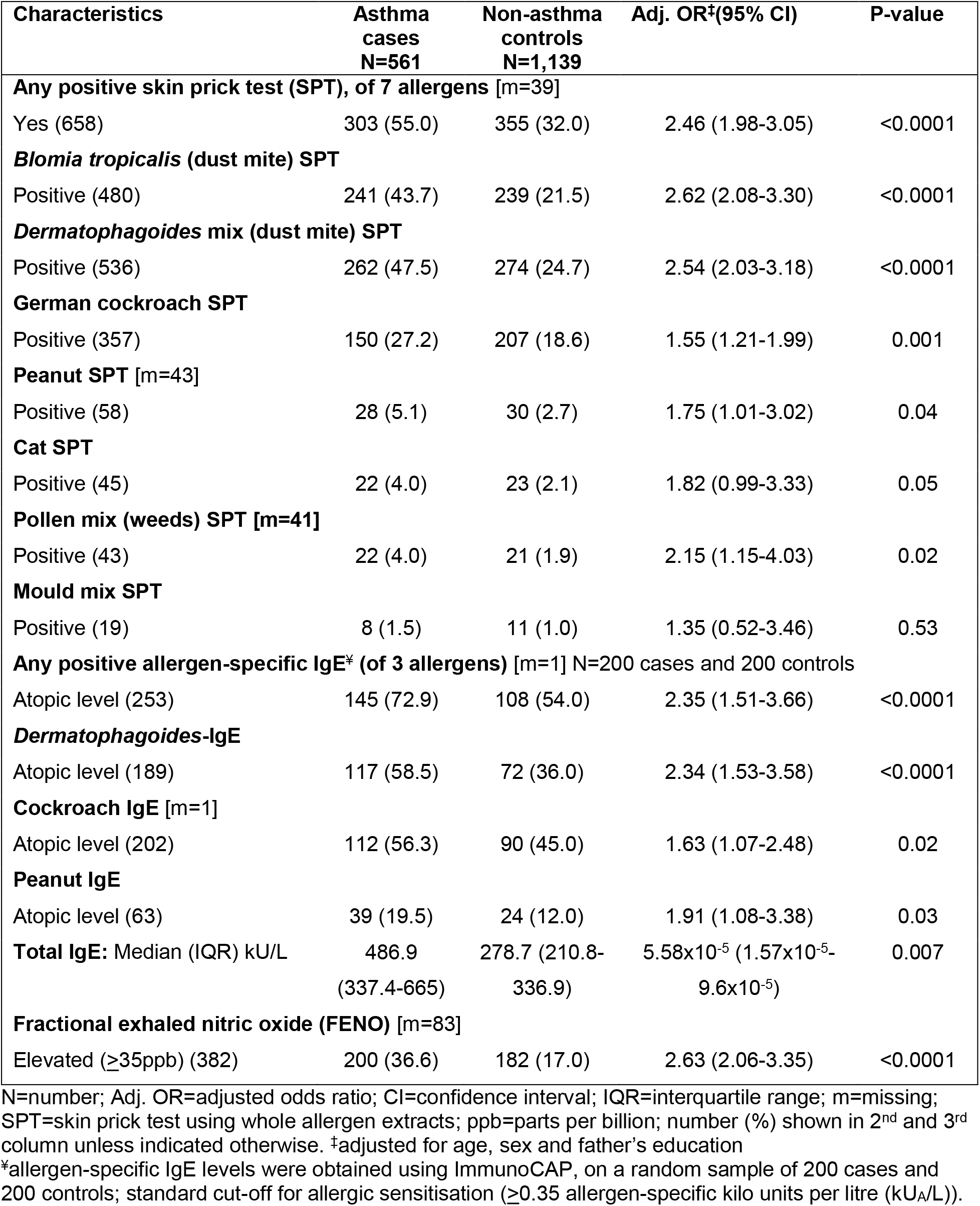
Atopy among asthma cases and non-asthma controls.

### Assessment of different combinations of risk factors for asthma

We investigated the relative importance of area of residence at birth versus the first years of life, on the asthma risk, by looking at children who migrated between rural and urban areas during these two periods. Children born and raised in rural areas had the lowest risk (the reference group); children born and raised in urban areas had the highest risk [2.37 (1.69-3.33)], children born in the urban area who migrated and spent most of 0-5 years in rural areas still had an increased risk of asthma [2.07 (1.14-3.77)], unlike children who were born in rural areas but migrated and spent most of 0-5 years in urban areas (Table 4).

**Table 4:** Area of residence at birth and in first five years of life, and asthma risk.

| Area of residence | Asthma cases<br>N=560 | Non-asthmatic controls<br>N=1,132 | Adj. OR<sup>‡</sup> (95% CI) | P-value |
| --- | --- | --- | --- | --- |
| Born and spent first 5 years in rural (267) | 50 (8.9) | 217 (19.2) | 1 | <0.0001 |
| Born in rural, spent first 5 years in urban (88) | 25 (4.5) | 63 (5.6) | 1.53 (0.86-2.71) |  |
| Born in urban, spent first 5 years in rural (72) | 22 (3.9) | 50 (4.4) | 2.07 (1.14-3.77) |  |
| Born and spent first 5 years in urban (1,265) | 463 (82.7) | 802 (70.8) | 2.37 (1.69-3.33) |  |

Adj. OR=adjusted odds ratio; CI=confidence interval; N=number; number (%) shown in 2<sup>nd</sup> and 3<sup>rd</sup> column
<sup>‡</sup>Adjusted for age, sex, and father's education

We investigated the combined effects of the child’s area of residence at birth and parental history of allergic disease. Even among children with no parental history of allergic disease, compared to being born in a rural area, being born in a town [2.01 (1.27-3.39)] or in the city [3.32 (1.64-6.74)] was associated with an increased risk of asthma, Table 5. Compared to the same reference group, children with a parental history of allergic disease had a higher risk of asthma that increased steadily from among children born in rural areas [2.77 (1.59-4.84)], in a town [4.68 (2.99-7.34)] and in the city [5.31 (2.94-9.62)]; p<0.0001 (Table 5). The relationship between children’s area of residence at birth and parental history of allergy was additive (interaction p-value=0.48).

**Table 5:**
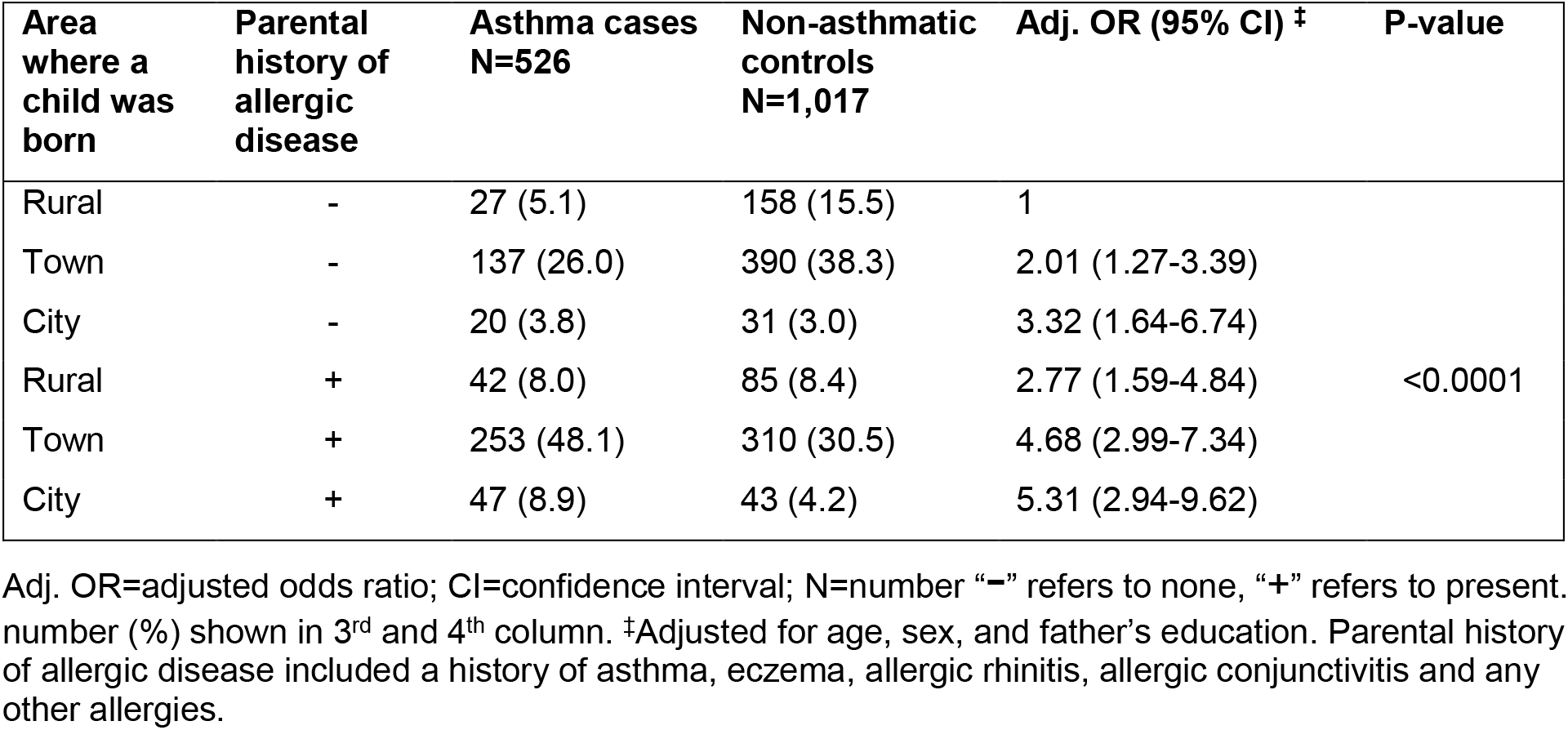
Area of residence at birth, parental history of allergic disease, and risk of asthma.

We also investigated the combined effects of area of residence at birth and children’s atopic status (positive SPT to any of seven allergens). Taking non-atopic children born in rural areas as the reference group, we found that non-atopic children born in town had a modest increase in asthma risk [1.60 (1.09-2.35)], which increased further among city-born children [2.50 (1.37-4.58)], Table 6. However, atopic children had a modestly increased risk of asthma even if they were born in the rural area [1.78 (1.04-3.05)], which increased substantially among atopic children born in town [4.17 (2.82-6.16)] or in the city [4.84 (2.72-8.59], Table 6. This relationship between area of residence and children’s atopy was additive (interaction p-value=0.38).

**Table 6:** Area of residence at birth, atopy status and risk of asthma.

| Area where a child was born | Atopy (SPT to any of 7 allergens) | Asthma cases<br>N=551 | Non-asthmatic controls<br>N=1,106 | Adj. OR (95% CI) ‡ | P-value |
| --- | --- | --- | --- | --- | --- |
| Rural | - | 43 (7.8) | 192 (17.4) | 1 | <0.0001 |
| Town | - | 180 (32.7) | 517 (46.7) | 1.60 (1.09-2.35) |  |
| City | - | 25 (4.5) | 43 (3.9) | 2.50 (1.37-4.58) |  |
| Rural | + | 31 (5.6) | 82 (7.4) | 1.78 (1.04-3.05) |  |
| Town | + | 229 (41.6) | 238 (21.5) | 4.17 (2.82-6.16) |  |
| City | + | 43 (7.8) | 34 (3.1) | 4.84 (2.72-8.59) |  |

Adj. OR=adjusted odds ratio; CI=confidence interval; N=number “-” refers to none, “+” refers to present; SPT-skin prick test to crude extracts of *Blomia tropicalis*, Dermatophagoides mix, cockroach, peanut, cat, weeds pollen mix, and mould mix. number (%) shown in 3<sup>rd</sup> and 4<sup>th</sup> column. †Adjusted for age, sex and father’s education.

In order to investigate the combined effects of parental education and urbanicity on asthma risk, we looked at the father’s education level in combination with the child’s area of residence at birth. We found that for each level of father’s education, there was an increasing risk gradient from rural-town-city; for each area of residence, increasing father’s education level was associated with increased asthma risk; the highest risk of asthma was among children born in the city whose fathers had a tertiary education [7.02 (3.69-13.37)] (Table 7). The relationship between father’s education and child’s area of residence at birth was additive (interaction p-value=0.17). The same pattern in Table 7 was seen for mother’s education and for child’s area of residence in the first five years of life.

**Table 7:**
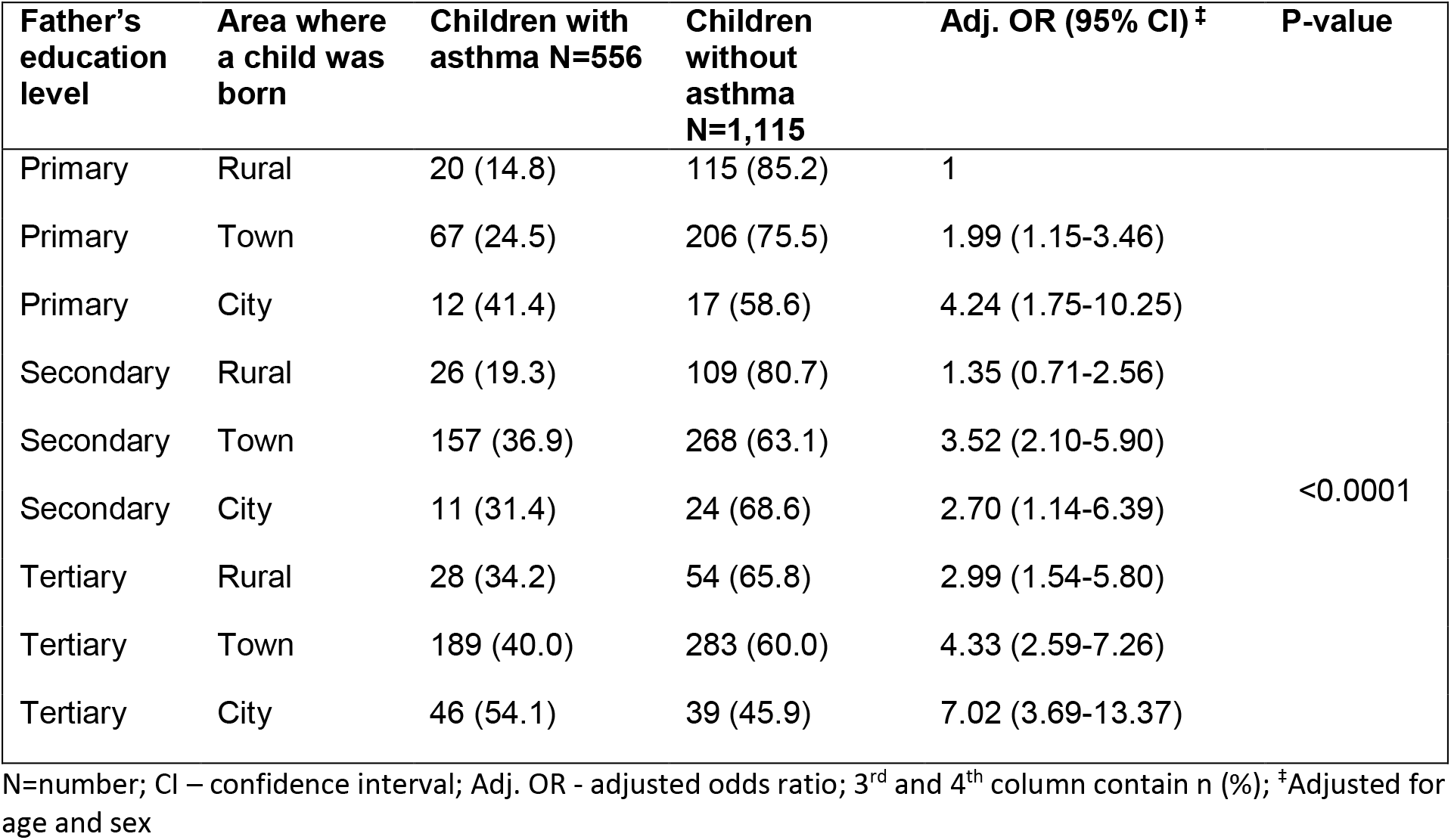
Father’s education, child’s area of residence at birth and the risk of asthma (N=1,671)

For the combined effect of atopy (SPT positive) and parental history of allergic disease, we found that asthma risk almost doubled among children with both atopy and a parental history of allergic disease [5.55 (3.98-7.75)] (Supplementary Table 2). However, children who had both parents with a history of allergic disease had an effect size similar to children who had either parent with a history of allergic disease [2.48 (1.75-3.51)] (Supplementary Table 3).

## Discussion

We provide novel findings that among schoolchildren in an urban area in Uganda, one of the main early-life predictors for asthma was the child’s area of residence at the time of birth and in their first five years of life. Children born and raised in rural areas had the lowest risk, children born and raised in small towns had a 2-fold increase in risk, while children born and raised in the city had a 3-fold increase in asthma risk. This is the first study in Africa to show such a strong gradient in asthma risk by place of birth. Previous studies have been mostly cross-sectional and conducted in either rural or urban settings, and therefore focusing on current residence; these have shown that the prevalence of asthma is lower among rural residents compared to urban residents(6, 7, 10).

Our findings confirm that the area where a child is born (usually the same as the area where the mother was resident during pregnancy) is important for asthma risk. Indeed, we found that when children moved to other environments, they carried with them the asthma risk related to their area of residence in pregnancy: children born in urban areas who were subsequently raised in rural areas still had a 2-fold increase in asthma risk, similar to their counterparts born and raised in urban areas. These findings are comparable to observations from Europe showing that children born on a farm (usually in rural areas) have a lower risk of asthma in later life, even when they subsequently moved to urban areas(27), and that children who migrated to Europe after age five had a lower prevalence of asthma, similar to their country of origin, than children who were either born in Europe or migrated before the age of five (28, 29). However, there are also studies that show increased risk of wheezing and allergy among children following migration(30, 31). Unlike studies from Europe and North America which have reported a lower risk of asthma for children born or raised on farms(32, 33), our study found no association between asthma risk and exposure to farm animals either during pregnancy or in early life. We hypothesise that this may be due to ubiquitous farm animal exposure due to subsistence farming, a widespread practice in Uganda, even in towns.

Our observation that children with asthma were more likely to have parents with tertiary education and to use gas or electricity for indoor cooking (as opposed to charcoal stoves) has been made by an earlier study in Uganda(14). Similarly, our finding that children with asthma reported the highest frequency of ‘trucks passing on the street near their home’ has been reported elsewhere(34–36). We suggest that these factors are proxy measures of a higher socio-economic status of asthma cases and of urbanicity, consistent with findings from other LMICs that have found a higher prevalence of asthma among children (and adults) in urban than rural areas(7, 37, 38). However, our findings contradict those from HICs in which asthma is associated with lower parental education(39) and socio-economic status(40). This suggests that there may be similarities in lifestyle and environmental factors between the highly educated and high socio-economic status families in LMICs with the low educated and low socio-economic status families in HICs, which increase asthma risk, and therefore require further investigation.

Although children with asthma were more likely to be sensitised to allergens than controls, the pattern of allergic sensitisation was similar among cases and controls; majorly sensitised to house-dust mites and cockroach, and least sensitised to peanut, cat, pollen and mould. This SPT response pattern was similar to other studies from Africa(15, 41), but different from Europe where the main allergens are dust mite, cat and pollen(42).

Although maternal smoking is a known risk factor for childhood asthma(43), our study found no association between maternal smoking and asthma. We attribute this to the low prevalence of smoking in this population, by the mothers during pregnancy (2.6%) and by any household members currently (<12%). The lack of association between asthma and indoor cooking with biomass fuels in this study is consistent with other studies from Africa(17, 18). Indeed, this is consistent with the general pattern of lower risk of asthma in rural areas (where biomass fuel use is highest) than urban areas.

We found an inverse association between asthma and current infection with any helminths, particularly *T. trichuria*. However, the association between asthma and current infections in Africa has been inconsistent across studies(44). What was novel in this study is that we also collected data on history of using de-worming medication in the last 12 months. We found that children with asthma were more likely to have used de-worming medication more than twice in the last 12 months compared to controls, and this was de-worming with albendazole (for geo-helminths) not with praziquantel (for schistosomiasis). This implies that the inverse association we noted between asthma and helminths may be partly explained by increased de-worming among children with asthma. The current de-worming schedule in this age-group in Uganda includes mass-deworming in schools, bi-annually for albendazole and once a year for praziquantel. We did not establish whether the additional de-worming was self-medicated or prescribed by medical workers.

We found that children with asthma were less likely to have a BCG scar, but there was no association with tuberculin skin test at enrolment. In Uganda, BCG vaccination is routinely given at birth. A lack of BCG scar does not mean the vaccine was not administered, but may indicate differences in immune responses among children who will eventually develop asthma. Alternatively, BCG vaccination may be protective against asthma, as suggested by previous studies(45).

The other important risk factors for asthma, that have been previously described, included parental history of allergic disease(46), a child’s atopy status and concomitant other allergic disease(47). The strength of this study was to demonstrate that having a combination of any two of these known risk factors for asthma had an additive effect of asthma risk, and that this risk also increased in relation to area of residence in early life, with an increasing rural-town-city gradient. This gradient and independent effects of parental education provide strong evidence for the role of environmental and lifestyle factors associated with urbanisation that are responsible for the increasing asthma risk in urban areas. More investigations are required to identify the specific factors, in order to design interventions to modify them so as to prevent the establishment of asthma risk in early life.

This study had limitations inherent to all case-control studies, such as potential recall bias, selection bias and confounding. We minimised recall bias by focusing on major early life events that a parent was likely to remember. It was re-assuring to note a strong correlation between the recalled events and objective measures such as SPT. We minimised the selection bias for controls by randomly selecting controls from the same class register where the cases were obtained, and by randomly selecting the 400 participants for the asIgE assay. We minimised confounding by adjusting for measured confounders in all our analyses, but cannot rule out the possible role of unmeasured confounders. Lastly, a large number of children were not included in this study, mostly because they were in the boarding school section and were unable to contact their parents/guardians in time to attend the parents’ meeting (to provide consent). Children in the boarding section were more likely to reside outside the study area, nonetheless, this may have introduced a selection bias.

## Conclusion

The risk of asthma among schoolchildren in urban Uganda is strongly predicted by their area of residence in early life, particularly at birth, with the highest risk among children whose early life is spent in small towns and in the city. This risk increases further in the presence of other asthma risk factors such as parental history of allergic disease, children’s own atopy, higher measures of socio-economic status and urbanicity. Given the current rapid urbanisation in Africa and other LMICs, the prevalence of asthma is likely to increase further. This study provides the basis for future studies investigating environmental and lifestyle factors that increase asthma risk in the urban areas of LMICs.

## Acknowledgements

We thank the study participants, their parents and guardians for their enthusiastic participation. We thank the teachers and school administrations for providing us with a conducive environment in which to conduct this study. We acknowledge the support we received from the Entebbe Municipality and Wakiso District Education Officials. Many thanks to Kisubi Hospital staff, particularly Dr. Robert Asaba and Dr. Rogers Sendijja, for their support during the results dissemination meetings we held in their premises. Many thanks to Ronald van Ree from Leiden University Medical Centre (The Netherlands) for working with GN on the asIgE ImmunoCAP assays.

---

**Supplementary Table 1:** Helminth infections and the risk of asthma.

| Type of helminth infection<sup>a</sup> | Asthma cases<br>N=518 | Non-asthma controls<br>N=1,048 | Adj. OR<sup>‡</sup> (95% CI) | P-value |
| --- | --- | --- | --- | --- |
| **Any helminth infection** |  |  |  |  |
| None (1,353) | 463 (89.4) | 890 (84.9) | 1 |  |
| Yes (213) | 55 (10.6) | 158 (15.1) | 0.72 (0.51-1.02) | 0.06 |
| ***Schistosoma mansoni*** |  |  |  |  |
| None (1,434) | 481 (92.9) | 953 (90.9) | 1 |  |
| Positive (132) | 37 (7.1) | 95 (9.1) | 0.78 (0.52-1.18) | 0.25 |
| ***Trichuris trichiura*** |  |  |  |  |
| None (1,526) | 512 (98.8) | 1,014 (96.8) | 1 |  |
| Yes (40) | 6 (1.2) | 34 (3.2) | 0.32 (0.12-0.85) | 0.02 |
| **Hookworm** |  |  |  |  |
| None (1,547) | 515 (99.4) | 1,032 (98.5) | 1 |  |
| Yes (19) | 3 (0.6) | 16 (1.5) | 0.45 (0.13-1.61) | 0.22 |
| ***Ascaris lumbricoides*** |  |  |  |  |
| None (1, 560) | 517 (99.8) | 1,043 (99.5) | 1 |  |
| Yes (6) | 1 (0.2) | 5 (0.5) | 0.57 (0.06-5.05) | 0.62 |
| **Other helminth infections<sup>a</sup>** |  |  |  |  |
| None (1,529) | 506 (97.7) | 1,023 (97.6) | 1 |  |
| Yes (37) | 12 (2.3) | 25 (2.4) | 1.34 (0.65-2.78) | 0.43 |

N=number; Adj. OR=adjusted odds ratio; CI=confidence interval; number (%) shown in 2<sup>nd</sup> and 3<sup>rd</sup> column
<sup>a</sup> All helminths infections based on Kato Katz analysis of three stool samples
<sup>‡</sup>Adjusted for child's age, sex, area of residence at birth, father's education and reported worm treatment in the last 12 months.
<sup>‡</sup>Other helminth infections included *Hymenolepis nana* and *Enterobius vermicularis*

**Supplementary Table 2:** Atopy status, parental history of allergic disease, and risk of asthma.

| **Atopy**<br>(SPT to<br>any of 7<br>allergens) | **Parental**<br>**history of**<br>**allergic**<br>**disease** | **Asthma**<br>**cases**<br>**N=518** | **Non-**<br>**asthmatic**<br>**controls**<br>**N=994** | **Adj. OR (95% CI) <sup>‡</sup>** | **P-value** |
| --- | --- | --- | --- | --- | --- |
| - | - | 77 (14.8) | 401 (40.3) | 1 |  |
| + | - | 105 (20.3) | 168 (16.9) | 2.98 (2.10-4.25) | <0.0001 |
| - | + | 149 (28.8) | 275 (27.7) | 2.69 (1.95-3.70) | <0.0001 |
| + | + | 187 (36.1) | 150 (15.1) | 5.55 (3.98-7.75) | <0.0001 |

Adj. OR=adjusted odds ratio; CI=confidence interval; N=number “-” refers to none, “+” refers to present. number (%) shown in 3<sup>rd</sup> and 4<sup>th</sup> column. <sup>‡</sup>Adjusted for age, sex, father’s education and area of residence where child was born. Parental history of allergic disease included a history of asthma, eczema, allergic rhinitis, allergic conjunctivitis and any other allergies.

**Supplementary Table 3:** Maternal and paternal history of allergic disease and the risk of asthma among children.

| **Parental reported history of**<br>**allergic disease** | **Asthma**<br>**cases**<br>**N=511** | **Non-**<br>**asthmatic**<br>**controls**<br>**N=996** | **Adj. OR<sup>‡</sup> (95%**<br>**CI)** | **P-value** |
| --- | --- | --- | --- | --- |
| Neither parents had history of<br>allergic disease | 184 (36.0) | 579 (58.1) | 1 |  |
| Only mother had history of<br>allergic disease | 159 (31.1) | 234 (23.5) | 2.06 (1.58-2.69) |  |
| Only father had history of allergic disease | 89 (17.4) | 87 (8.7) | 2.69 (1.90-3.80) |  |
| Both parents had history of allergic disease | 79 (15.5) | 96 (9.6) | 2.48 (1.75-3.51) | <0.0001 |

Adj. OR=adjusted odds ratio; CI=confidence interval; N=number; number (%) shown in 2<sup>nd</sup> and 3<sup>rd</sup>column.
‡Adjusted for age, sex, father's education and area of residence at the time of birth
Parental history of allergic disease included a history of asthma, eczema, allergic rhinitis, allergic conjunctivitis and any other allergies.

## References

1. WHO. Asthma Fact Sheet. 2017.

2. Asher MI, Montefort S, Bjorksten B, Lai CK, Strachan DP, Weiland SK, et al. Worldwide time trends in the prevalence of symptoms of asthma, allergic rhinoconjunctivitis, and eczema in childhood: ISAAC Phases One and Three repeat multicountry cross-sectional surveys. Lancet (London, England). 2006;368(9537):733–43.

3. Addo-Yobo EO, Woodcock A, Allotey A, Baffoe-Bonnie B, Strachan D, Custovic A. Exercise-induced bronchospasm and atopy in Ghana: two surveys ten years apart. PLoS Med. 2007;4(2):e70.

4. van Gemert F, van der Molen T, Jones R, Chavannes N. The impact of asthma and COPD in sub-Saharan Africa. Primary care respiratory journal: journal of the General Practice Airways Group. 2011;20(3):240–8.

5. Zar HJ, Ehrlich RI, Workman L, Weinberg EG. The changing prevalence of asthma, allergic rhinitis and atopic eczema in African adolescents from 1995 to 2002. Pediatric allergy and immunology: official publication of the European Society of Pediatric Allergy and Immunology. 2007;18(7):560–5.

6. Lawson JA, Rennie DC, Cockcroft DW, Dyck R, Afanasieva A, Oluwole O, et al. Childhood asthma, asthma severity indicators, and related conditions along an urban-rural gradient: a cross-sectional study. BMC pulmonary medicine. 2017;17(1):4.

7. Morgan BW, Siddharthan T, Grigsby MR, Pollard SL, Kalyesubula R, Wise RA, et al. Asthma and Allergic Disorders in Uganda: A Population-Based Study Across Urban and Rural Settings. The journal of allergy and clinical immunology In practice. 2018.

8. Lotvall J, Akdis CA, Bacharier LB, Bjermer L, Casale TB, Custovic A, et al. Asthma endotypes: a new approach to classification of disease entities within the asthma syndrome. The Journal of allergy and clinical immunology. 2011;127(2):355–60.

9. Douwes J, Pearce N. Asthma and the westernization ‘package’. International journal of epidemiology. 2002;31(6):1098–102.

10. Botha M, Basera W, Facey-Thomas HE, Gaunt B, Genuneit J. Nutrition and allergic diseases in urban and rural communities from the South African Food Allergy cohort. 2019.

11. Addo-Yobo EO, Custovic A, Taggart SC, Craven M, Bonnie B, Woodcock A. Risk factors for asthma in urban Ghana. The Journal of allergy and clinical immunology. 2001;108(3):363–8.

12. Ayuk AC, Ramjith J, Zar HJ. Environmental risk factors for asthma in 13-14 year old African children. Pediatric pulmonology. 2018;53(11):1475–84.

13. Arrais M, Lulua O, Quifica F, Rosado-Pinto J, Gama JMR, Taborda-Barata L. Prevalence of asthma, allergic rhinitis and eczema in 6-7-year-old schoolchildren from Luanda, Angola. Allergologia et immunopathologia. 2019.

14. Nantanda R, Ostergaard MS, Ndeezi G, Tumwine JK. Factors associated with asthma among under-fives in Mulago hospital, Kampala Uganda: a cross sectional study. BMC pediatrics. 2013;13:141.

15. Nyembue TD, Jorissen M, Hellings PW, Muyunga C, Kayembe JM. Prevalence and determinants of allergic diseases in a Congolese population. International forum of allergy & rhinology. 2012;2(4):285–93.

16. Mehanna N, Mohamed N. Allergy-related disorders (ARDs) among Ethiopian primary school-aged children: Prevalence and associated risk factors. 2018;13(9):e0204521.

17. Thacher JD, Emmelin A, Madaki AJ, Thacher TD. Biomass fuel use and the risk of asthma in Nigerian children. Respiratory medicine. 2013;107(12):1845–51.

18. Oluwole O, Arinola GO, Huo D, Olopade CO. Household biomass fuel use, asthma symptoms severity, and asthma underdiagnosis in rural schoolchildren in Nigeria: a cross-sectional observational study. BMC pulmonary medicine. 2017;17(1):3.

19. Oluwole O, Arinola GO, Huo D, Olopade CO. Biomass fuel exposure and asthma symptoms among rural school children in Nigeria. The Journal of asthma : official journal of the Association for the Care of Asthma. 2017;54(4):347–56.

20. von Elm E, Altman DG, Egger M, Pocock SJ, Gøtzsche PC, Vandenbroucke JP. The Strengthening the Reporting of Observational Studies in Epidemiology (STROBE) Statement: Guidelines for reporting observational studies. International Journal of Surgery. 2014;12(12):1495–9.

21. Asher MI, Keil U, Anderson HR, Beasley R, Crane J, Martinez F, et al. International Study of Asthma and Allergies in Childhood (ISAAC): rationale and methods. The European respiratory journal. 1995;8(3):483–91.

22. Mpairwe H, Muhangi L, Ndibazza J, Tumusiime J, Muwanga M, Rodrigues LC, et al. Skin prick test reactivity to common allergens among women in Entebbe, Uganda. Transactions of the Royal Society of Tropical Medicine and Hygiene. 2008;102(4):367–73.

23. Katz N, Chaves A, Pellegrino J. A simple device for quantitative stool thick-smear technique in Schistosomiasis mansoni. Rev Inst Med Trop Sao Paulo. 1972;14(6):397–400.

24. Nkurunungi G, Lutangira JE, Lule SA, Akurut H, Kizindo R, Fitchett JR, et al. Determining Mycobacterium tuberculosis infection among BCG-immunised Ugandan children by T-SPOT.TB and tuberculin skin testing. PLoS ONE. 2012;7(10):e47340.

25. Greenland S, Pearce N. Statistical foundations for model-based adjustments. Annual review of public health. 2015;36:89–108.

26. Greenland S, Daniel R, Pearce N. Outcome modelling strategies in epidemiology: traditional methods and basic alternatives. International journal of epidemiology. 2016;45(2):565–75.

27. Leynaert B, Neukirch C, Jarvis D, Chinn S, Burney P, Neukirch F. Does Living on a Farm during Childhood Protect against Asthma, Allergic Rhinitis, and Atopy in Adulthood? 2001;164(10):1829–34.

28. Migliore E, Pearce N, Bugiani M, Galletti G, Biggeri A, Bisanti L, et al. Prevalence of respiratory symptoms in migrant children to Italy: the results of SIDRIA-2 study. Allergy. 2007;62(3):293–300.

29. Kuehni CE, Strippoli MP, Low N, Silverman M. Asthma in young south Asian women living in the United Kingdom: the importance of early life. Clinical and experimental allergy : journal of the British Society for Allergy and Clinical Immunology. 2007;37(1):47–53.

30. Rodriguez A, Vaca MG, Chico ME, Rodrigues LC, Barreto ML, Cooper PJ. Rural to urban migration is associated with increased prevalence of childhood wheeze in a Latin-American city. BMJ open respiratory research. 2017;4(1):e000205.

31. Stein M, Greenberg Z, Boaz M, Handzel ZT, Meshesha MK, Bentwich Z. The Role of Helminth Infection and Environment in the Development of Allergy: A Prospective Study of Newly-Arrived Ethiopian Immigrants in Israel. PLoS neglected tropical diseases. 2016;10(1):e0004208.

32. Genuneit J. Exposure to farming environments in childhood and asthma and wheeze in rural populations: a systematic review with meta-analysis. Pediatric allergy and immunology: official publication of the European Society of Pediatric Allergy and Immunology. 2012;23(6):509–18.

33. Timm S, Frydenberg M, Janson C, Campbell B, Forsberg B, Gislason T, et al. The Urban-Rural Gradient In Asthma: A Population-Based Study in Northern Europe. International journal of environmental research and public health. 2015;13(1).

34. Sharma SK, Banga A. Prevalence and risk factors for wheezing in children from rural areas of north India. Allergy and asthma proceedings. 2007;28(6):647–53.

35. Shirinde J, Wichmann J, Voyi K. Association between wheeze and selected air pollution sources in an air pollution priority area in South Africa: a cross-sectional study. Environmental health: a global access science source. 2014;13(1):32.

36. Venn A, Yemaneberhan H, Lewis S, Parry E, Britton J. Proximity of the home to roads and the risk of wheeze in an Ethiopian population. Occupational and environmental medicine. 2005;62(6):376–80.

37. Gaviola C, Miele CH, Wise RA, Gilman RH, Jaganath D, Miranda JJ, et al. Urbanisation but not biomass fuel smoke exposure is associated with asthma prevalence in four resource-limited settings. Thorax. 2016;71(2):154–60.

38. Robinson CL, Baumann LM, Romero K, Combe JM, Gomez A, Gilman RH, et al. Effect of urbanisation on asthma, allergy and airways inflammation in a developing country setting. Thorax. 2011;66(12):1051–7.

39. Lewis KM, Ruiz M, Goldblatt P, Morrison J, Porta D, Forastiere F, et al. Mother’s education and offspring asthma risk in 10 European cohort studies. European journal of epidemiology. 2017;32(9):797–805.

40. Akinbami LJ, Moorman JE, Liu X. Asthma prevalence, health care use, and mortality: United States, 2005-2009. National health statistics reports. 2011(32):1–14.

41. Mbatchou Ngahane BH, Noah D, Nganda Motto M, Mapoure Njankouo Y, Njock LR. Sensitization to common aeroallergens in a population of young adults in a sub-Saharan Africa setting: a cross-sectional study. Allergy, Asthma, and Clinical Immunology: Official Journal of the Canadian Society of Allergy and Clinical Immunology. 2016;12:1.

42. Bousquet PJ, Chinn S, Janson C, Kogevinas M, Burney P, Jarvis D. Geographical variation in the prevalence of positive skin tests to environmental aeroallergens in the European Community Respiratory Health Survey I. Allergy. 2007;62(3):301–9.

43. Silvestri M, Franchi S, Pistorio A, Petecchia L, Rusconi F. Smoke exposure, wheezing, and asthma development: a systematic review and meta-analysis in unselected birth cohorts. Pediatric pulmonology. 2015;50(4):353–62.

44. Mpairwe H, Amoah AS. Parasites and allergy: Observations from Africa. 2019;41(6):e12589.

45. El-Zein M, Parent ME, Benedetti A, Rousseau MC. Does BCG vaccination protect against the development of childhood asthma? A systematic review and meta-analysis of epidemiological studies. International journal of epidemiology. 2010;39(2):469–86.

46. Lim RH, Kobzik L, Dahl M. Risk for asthma in offspring of asthmatic mothers versus fathers: a meta-analysis. PLoS ONE. 2010;5(4):e10134.

47. Bao Y, Chen Z, Liu E, Xiang L, Zhao D, Hong J. Risk Factors in Preschool Children for Predicting Asthma During the Preschool Age and the Early School Age: a Systematic Review and Meta-Analysis. Current allergy and asthma reports. 2017;17(12):85.

